# ComplexMatrixComb: Predicting Drug Combination IC_50_ Doses via Complex Numbers and Matrix Factorization

**DOI:** 10.1101/2025.10.06.680642

**Authors:** Mohammad Abdollahi, Asiyeh Mirzaei Koli, Shokoofeh Ghiam, Changiz Eslahchi

## Abstract

Determining precise drug concentrations to inhibit cancer cell growth is a critical but resource-intensive challenge, especially for combinations requiring many dose pairs. Existing computational methods often predict synergy or classify interactions but rarely estimate exact concentration pairs for a defined inhibitory effect. We introduce ComplexMatrixComb, a matrix factorization framework predicting drug combination dosages achieving IC_50_ for specific cancer cell lines. It encodes drug concentrations as real and imaginary components of complex numbers, modeling joint dose–response dynamics and enabling accurate IC_50_ estimation. Across O’Neil, NCI-ALMANAC, and AZ-Dream datasets, ComplexMatrixComb outperformed traditional machine learning in regression and classification, showing robustness to drug order and adaptability to varied designs. Integration with models such as ComboFM and ComboLTR showed predicted concentrations closely matched ground-truth doses. Experimental validation of five drug pair–cell line predictions using MTT assays confirmed 50% growth inhibition. By reducing reliance on exhaustive screens, ComplexMatrixComb offers a scalable, data-driven tool to accelerate preclinical research and support personalized oncology strategies.

## 1 Introduction

Cancer remains one of the leading health challenges worldwide, with millions of new cases diagnosed every year and high mortality rates in diverse populations. Numerous biological, genetic, and environmental factors contribute to the development and progression of different types of cancer [1–5]. In recent years, bioinformatics has emerged as a crucial discipline to manage the complexity of cancer research. It supports various aspects of drug discovery and development, including protein targeting studies, next-generation sequencing analysis, functional annotation of genes and proteins, and identification of therapeutic biomarkers [6–9]. The development of effective cancer therapies remains a major challenge in oncology, due to factors such as drug resistance, tumor heterogeneity, and the limited efficacy of single agents. One of the most widely used metrics for assessing drug potency is the half-maximal inhibitory concentration (IC_50_), which represents the concentration of a drug required to inhibit a biological or biochemical function by 50% [10]. Accurate estimation of IC_50_ plays a central role in drug discovery and pharmacology, as it enables researchers to evaluate and compare the potency of different substances in inhibiting cancer cell growth. In recent years, numerous studies have focused on improving IC_50_ prediction, particularly for single drugs, by employing advanced machine learning and computational techniques.

Recent advancements in IC_50_ prediction highlight the growing application of advanced machine learning and computational techniques to enhance precision in drug response modeling. For example, Devipriya and Vijaya (2024)[11] utilized a Graph Convolutional Neural Network (GCNN) to model IC_50_ values with higher precision by leveraging the structural properties of drug molecules and gene expression data relevant to neurodegenerative disorders. Their innovative approach, which incorporates graph representations and global features, outperforms traditional machine learning methods in accuracy. Similarly, Mosos et al. (2024)[12] integrated quantitative structure-activity relationships (QSAR) and structure-based drug design (SBDD) in a machine learning framework to predict IC_50_ values for Non-Nucleoside Reverse-Transcriptase Inhibitor (NNRTI) analogs, underscoring the potential of computational models in designing safer and more effective drugs.

Numerous other studies have explored IC_50_ prediction and drug sensitivity modeling using various approaches, such as machine learning, deep learning, graph theory, and recommender systems. For example, Emdadi and Eslahchi (2021)[13] approached the problem as a recommender system, employing methods like logistic matrix factorization and Auto-HMM-LMF, a feature selection-based approach combining autoencoders and hidden Markov models. In the realm of deep learning, Joo et al. (2019)[14] proposed DeepIC_50_, a deep learning model that leverages molecular fingerprints of drugs and mutation profiles to improve IC_50_ prediction accuracy across diverse cancer cell lines. Jiang et al. (2022)[15] introduced DeepTTA, a transformer-based model for predicting cancer drug response that combines drug chemical substructures and transcriptomic gene expression data. DeepTTA demonstrated superior performance compared to existing methods, making it an effective tool for anti-cancer drug design. Furthermore, Sagingalieva et al. (2023)[16] presented a novel hybrid quantum neural network architecture for drug response prediction, incorporating deep quantum circuits to improve prediction accuracy by 15% over classical methods. This innovative approach highlights the potential of quantum computing in advancing personalized medicine and cancer treatment strategies.

Graph theory and machine learning approaches have also been employed, as demonstrated by Yassaee Meybodi and Eslahchi (2021)[17], who identified optimal subsets of drugs to predict anti-cancer responses. Similarly, Ahmadi Moughari and Eslahchi introduced ADRML (2020)[18], a manifold learning-based approach to anticancer drug response prediction. Computational methodologies like the Density Functional Theory (DFT)-based approach by Bag (2021)[19] further showcase the potential of sophisticated computational models in addressing limitations inherent in traditional QSAR-based methods reliant on extensive datasets.

While assessing the potency of single drugs using metrics like IC_50_ is critical for understanding their therapeutic potential, cancer treatment often requires more than a single-agent approach. Drug combinations have emerged as a promising strategy to overcome challenges such as drug resistance, tumor heterogeneity, and limited efficacy of individual agents. By leveraging the complementary mechanisms of action between drugs, combination therapies aim to enhance treatment outcomes, reduce toxicity, and minimize the likelihood of resistance[20–22].

The study of drug combinations has focused on several key problems, including predicting drug synergy scores, classifying drug interactions, and predicting inhibition rates. Each of these problems addresses a different aspect of understanding and optimizing drug combinations.

Drug synergy occurs when the combined effect of two drugs is greater than the sum of their individual effects. Conversely, drug antagonism happens when the combined effect is less than the sum of their individual effects, leading to a reduced therapeutic outcome. Additivity refers to a situation where the combined effect of the drugs is exactly the sum of their individual effects, implying no interaction[23, 24]. Various methods have been developed to predict these interactions. For instance, deep learning models like DeepSynergy by Preuer et al. (2018)[25] and DeepDDS by Zhang et al. (2021)[26] leverage chemical structure and gene expression profiles to predict drug synergy. Additionally, the MARSY model by El Khili et al. (2023)[27] presents a multitask deep-learning framework that incorporates gene expression profiles of cancer cell lines, as well as the differential expression signature induced by each drug, to predict drug-pair synergy scores. Machine learning approaches such as Gradient Boosting Machines (GBMs) have also been utilized, with Liu et al. (2019)[28] employing GBMs in their DrugComb model to integrate multi-omics data and predict drug synergy. Xu et al. (2023)[29] introduced DFFNDDS, a deep learning model that uses a dual feature fusion mechanism and fine-tuned pre-trained language models to predict synergistic drug combinations, outperforming competitive methods and providing a reliable tool for identifying such combinations.

Classification of drug interactions involves determining whether a combination of drugs will result in synergistic, antagonistic, or additive effects. Methods in this area include SynergyFinder by Ianevski et al. (2020)[30], which offers an interactive analysis of drug combination screening results to predict synergy, antagonism, and additivity. Similarly, SynToxProfiler by Ianevski et al. (2020)[31] uses machine learning to classify drug interactions and predict toxicity outcomes. The MultiSyn model by Monem et al. (2024)[32] leverages multi-task learning with graph-based and SMILES-based drug features and integrates cancer cell line gene expression to classify drug synergy outcomes. Wang et al. (2023)[33] introduced DEML, an ensemble-based multi-task neural network, which simultaneously predicts drug synergy, DDI classification, and other synergy-related tasks, using a gating mechanism to enhance task-specific fusion. Lastly, SYNDEEP by Torkamannia et al. (2023)[34] applies deep neural network classification, combining genomic and interaction data, to achieve high accuracy in predicting cancer drug synergy.

Predicting the inhibition rate of drug pairs involves determining how effective a combination of drugs is in inhibiting cancer cell growth, which is crucial for understanding the practical impact of drug combinations in therapeutic settings. Notable efforts include ComboFM by Julkunen et al. (2020)[35], which uses multi-way interactions for systematic prediction of pre-clinical drug combination effects. PIICM, developed by Rønneberg et al. (2023)[36], applies a probabilistic framework to predict dose-response surfaces in high-throughput drug combination datasets, incorporating experimental uncertainty to capture relevant synergistic features. Similarly, comboLTR by Wang et al. (2021)[37] uses latent tensor reconstruction with polynomial regression to model drug combination effects efficiently across doses and cellular contexts, achieving high accuracy for new drug combinations.

Despite these advances, there remains a significant gap in research regarding the direct prediction of IC_50_ values for drug combinations on specific cancer cell lines. Most existing models and methodologies focus on synergy scores or interaction classification without providing a method to predict the precise concentrations needed for a 50% inhibition of cancer cell growth. This gap highlights the need for novel approaches that can accurately predict these concentrations, thereby providing more practical and actionable insights for combination therapies.

To address the critical and previously unaddressed challenge of predicting precise drug pair concentrations for cancer treatment, we present ComplexMatrixComb, a novel matrix factorization framework designed to estimate the exact concentrations of two drugs required to achieve 50% inhibition (IC_50_) of specific cancer cell lines. In our approach, each drug’s concentration is encoded as either the real or imaginary component of a complex number, enabling the model to capture intricate, joint dose–response dynamics with high fidelity. Unlike conventional real-valued models, this complex-number formulation allows ComplexMatrixComb to model multidimensional interactions between drug combinations and cellular responses more effectively. By filling a significant gap in computational pharmacology, our method offers a transformative tool for optimizing combinatorial cancer therapies.

The term ComplexMatrix reflects our novel use of matrix factorization with complex numbers to model intricate drug–drug and drug pair–cell interactions. Unlike traditional real-valued approaches, this complex-number framework offers greater expressive power for capturing subtle, multidimensional relationships between drug combinations and their effects on cancer cells. By integrating advanced machine learning techniques, our method goes beyond synergy estimation to accurately predict the optimal concentrations required to achieve a targeted inhibitory response.

We evaluate ComplexMatrixComb on three benchmark datasets—O’Neil, NCIALMANAC, and AZ-DREAM—and show that it consistently outperforms traditional models in both regression and classification tasks. The model demonstrates robustness to drug ordering and generalizes well across heterogeneous experimental designs. Moreover, when integrated into existing frameworks such as ComboFM and ComboLTR, our predicted drug concentrations yield inhibition results that closely match those obtained using ground-truth dosing.

ComboFM is a machine learning framework that leverages higher-order tensor factorization to model context-specific drug interactions. It uses factorization machines to efficiently learn latent representations of drugs and cell lines, enabling accurate response prediction even in sparse experimental settings by transferring knowledge from similar combinations.

ComboLTR, in contrast, employs polynomial regression and latent tensor reconstruction to model complex, nonlinear relationships in drug response data. By combining recommender system–style indexing with chemical and multi-omics features, ComboLTR can infer full dose–response matrices for novel drug pairs. It excels at predicting combination effects across a range of concentrations and cellular contexts, achieving state-of-the-art accuracy with high computational efficiency.

To assess real-world applicability, we experimentally tested five high-confidence drug pair–cell line predictions using MTT assays. The results confirmed that our predicted concentrations consistently induced approximately 50% growth inhibition, closely matching IC_50_ expectations. By minimizing the need for exhaustive empirical screening, ComplexMatrixComb offers a scalable and practical solution for dose prediction in drug combination therapy, advancing both preclinical research and the development of personalized treatment strategies.

In the subsequent sections, we will thoroughly explain our proposed method and the datasets utilized in this study. We will then present our evaluation metrics and results, providing a comprehensive analysis of our model’s performance. Specifically, we selected four drug pairs from the ALMANAC dataset and four from the O’Neil dataset. Using the predicted concentrations, we performed laboratory experiments on these previously untested drug-cell line combinations to evaluate the real-world applicability of our approach. The details and findings of this case study will be discussed in the case study section. Finally, in the discussion section, we will offer a deeper exploration of our method, its implications, and potential future directions for research.

## 2 Method & Materials

### 2.1 Dataset

In this project, we used the O’Neil dataset produced by Merck & Co. [38]. The original dataset contains 38 unique drugs; however, since one drug lacked any usable concentration–response data, it was excluded from our analysis. As a result, our study was conducted on 37 unique drugs. The dataset also consists of 583 unique drug pairs, each tested against 39 human cancer cell lines derived from 7 different tissue types. In total, the dataset comprises 368832 samples, where each sample consists of two drugs, a cell line, and the inhibition rate of the cell line after applying different concentration values of the two compounds.

Another dataset utilized in this study was derived from the NCI-ALMANAC [39], the largest publicly available drug combination dataset. The original dataset encompasses over 5,000 combinations of approximately 100 FDA-approved small molecule drugs screened against the NCI-60 panel of human tumor cell lines representing nine tissue types. It includes over 3 million response measurements across various concentrations. Similar to the ComboFM method, we adopted the same sampling approach to optimize computational efficiency. A subset of the dataset was created by randomly sampling 50 drugs (Supplementary Table 3) while maintaining the original distribution of drug combination responses. Drug combinations with complete measurements across all 60 cell lines were retained, resulting in a refined dataset containing 617 drug combinations of 50 unique drugs screened in 45 concentrations, yielding 333,180 response measurements.

Another dataset utilized in this project is the AstraZeneca-DREAM (AZ-DREAM) [40] challenge dataset, which provides a valuable resource for drug combination analysis. This dataset consists of experimental data on the effects of drug combinations across various cancer cell lines. It includes 91 unique drugs tested in 11,576 distinct combinations on 85 human cancer cell lines derived from diverse tissue types. Each combination is represented by the inhibition rate of the respective cell line under different drug concentration pairs, offering a comprehensive view of drug interaction effects. The AZ-DREAM dataset is widely recognized for its extensive coverage and is frequently used as a benchmark in drug synergy prediction studies.

It is worth emphasizing that these three datasets, Merck, NCI-ALMANAC, and AstraZeneca-DREAM, are the only publicly available resources that provide detailed information on drug combinations, including the inhibition rates of cell lines under various concentration pairs of two drugs, along with comprehensive coverage of tissue types. While other datasets, such as DrugComb, exist, they do not include concentration-specific inhibition data, making the three datasets used in this study uniquely essential for our analysis.

### 2.2 Method

Suppose *D* = {*d*_1_, *d*_2_, …, *d*_*n*_} and *C* = {*c*_1_, *c*_2_, …, *c*_*m*_} are the drugs and cell lines set in the dataset, respectively. Besides, for each pair of drugs (*d*_*i*_, *d*_*j*_) and cell line c, there is a k×k drug-response matrix whose entries show the cell line inhibition rate after applying different drug concentrations of these drug pairs. For instance, considering the drug pair (5-FU, BEZ-235) and the A2058 cell line from the O’Neil dataset, the data can be organized into a 4×4 matrix. The rows represent increasing concentrations of 5-FU, while the columns correspond to increasing concentrations of BEZ-235, with all drug concentrations reported in the micromolar (µM) scale. Each cell in the matrix reflects the observed inhibition rate for the respective combination of concentrations of A2058 (see Figure 1).

**Fig. 1.**
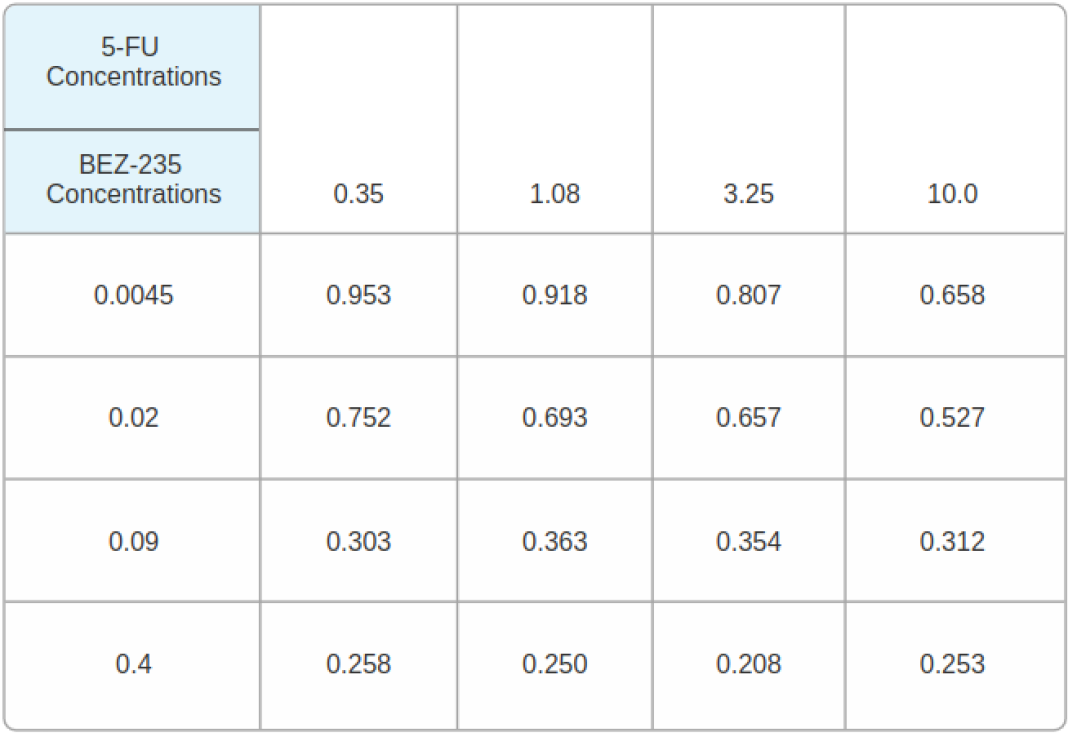
This figure illustrates a record from the O’Neil dataset, showing inhibition rates for the drug pair (5-FU, BEZ-235) applied to the A2058 cell line. The matrix represents drug concentrations in the micromolar (µM) scale, with rows corresponding to 5-FU concentrations and columns to BEZ235 concentrations.

The main goal in ComplexMatrixComb is to implement an algorithm that predicts drug concentrations *µ*_*i*_ and *µ*_*j*_ for a pair of drugs such as *d*_*i*_ and *d*_*j*_ that with these concentrations in a specific cell line, leads to inhibition rate close to 50 percent (between 45% and 55%).

The following are the steps of ComplexMatrixComb:

1. Scaling drug concentrations
2. IC_50_ selection set
3. Create cell line-drug matrix with complex entries
4. Matrix factorization method
5. Predict concentration values for new drug pairs

In the following, the steps of the method are explained in detail and Figure 2 demonstrates the workflow of our method.

**Fig. 2.**
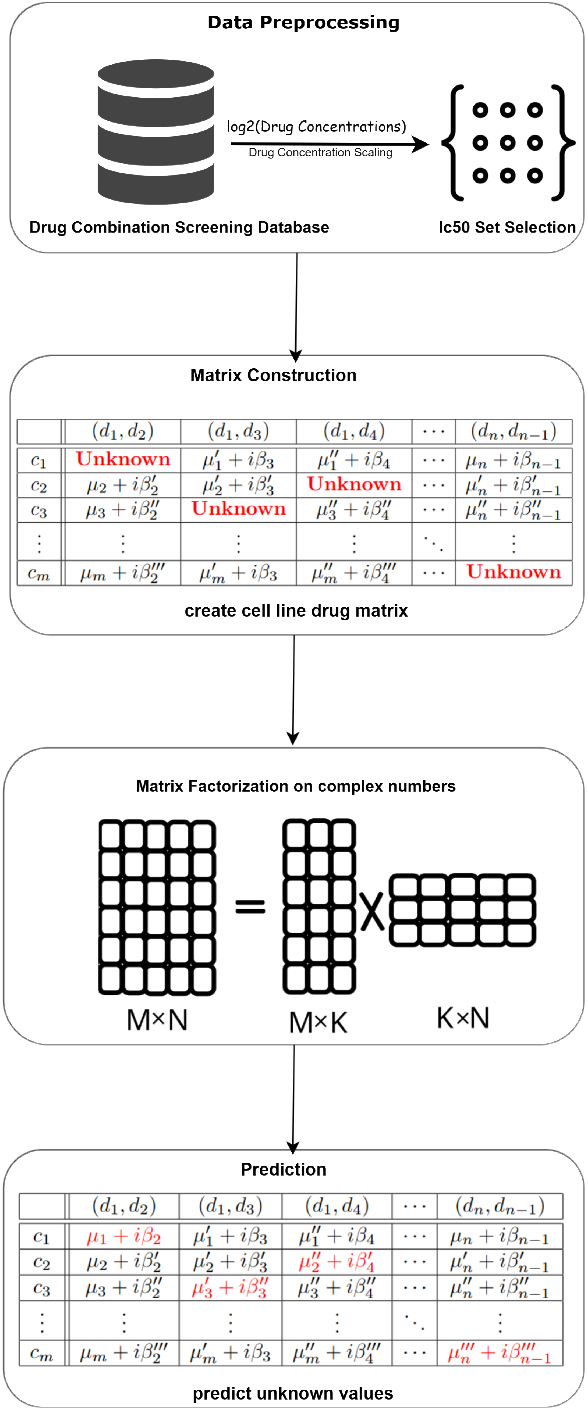
Schematic representation of the ComplexMatrixComb workflow. The pipeline includes preprocessing drug concentration matrices, selecting IC_50_ inhibition points, constructing a complex-valued drug-cell line matrix, applying modified matrix factorization, and predicting effective drug pair concentrations that yield approximately 50% inhibition for specific cell lines.

#### 2.2.1 Step1: Scaling drug concentrations

For each pair of drugs (*d*_*i*_, *d*_*j*_) and cell line c, there is a k×k matrix whose elements show the cell line inhibition rate after applying different drug concentrations of these drug pairs. The matrix size is 4×4 for the O’Neil dataset, 3×3 for the NCI-ALMANAC dataset, and 5×5 for the AZ-DREAM dataset. To ensure the convergence of our algorithm, it is essential to address the variability in drug concentration values. A wide range of concentration values can hinder the convergence of our algorithm, making it challenging to achieve consistent and reliable results. To mitigate this issue, we will transform the drug concentration values by taking the logarithm base 2 (*log*_2_) of each value. By doing so, we achieve a more normalized distribution of values, which facilitates better convergence and stability in our computational methods.

#### 2.2.2 Step2: IC_50_ set selection

Suppose S denotes the IC_50_ selection set, defined as follows: 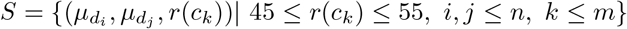 where 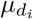 and 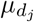 represent the concentrations for drugs *d*_*i*_ and *d*_*j*_, respectively, and *r*(*c*_*k*_) denotes the inhibition rate for cell line *c*_*k*_ after applying the drug pair (*d*_*i*_, *d*_*j*_) at these concentrations.

#### 2.2.3 Step3: Creating drug-cell line matrix with complex entries

In this step, we construct a matrix *M* = [*m*_*k*,(*i,j*)_] corresponding to the set S. Each entry *m*_*k*,(*i,j*)_ represents the value for column k, corresponding to cell line *c*_*k*_ and row (i,j) corresponding to the drug pair (*d*_*i*_, *d*_*j*_). This entry is defined as: 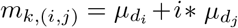, where 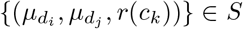. It is important to note that IC_50_ concentrations are not available for every drug pair across all cell lines, resulting in some missing values in the matrix. To address this, we remove columns (representing drug pairs) that contain more than 50% missing values. Nevertheless, some missing entries still remain in the resulting matrix *CD*. These missing values were never treated as real observations: in our implementation, they were masked during training and therefore did not contribute to the optimization process. For technical reasons in the released code, these missing entries appear as the placeholder value 0 + 0*j*; however, this representation was used solely as a marker and was ignored during model fitting. Thus, the model was trained exclusively on valid concentration–response data without introducing any artificial zero-valued samples. The schematic representation of this matrix is shown in Figure 3.

**Fig. 3.**
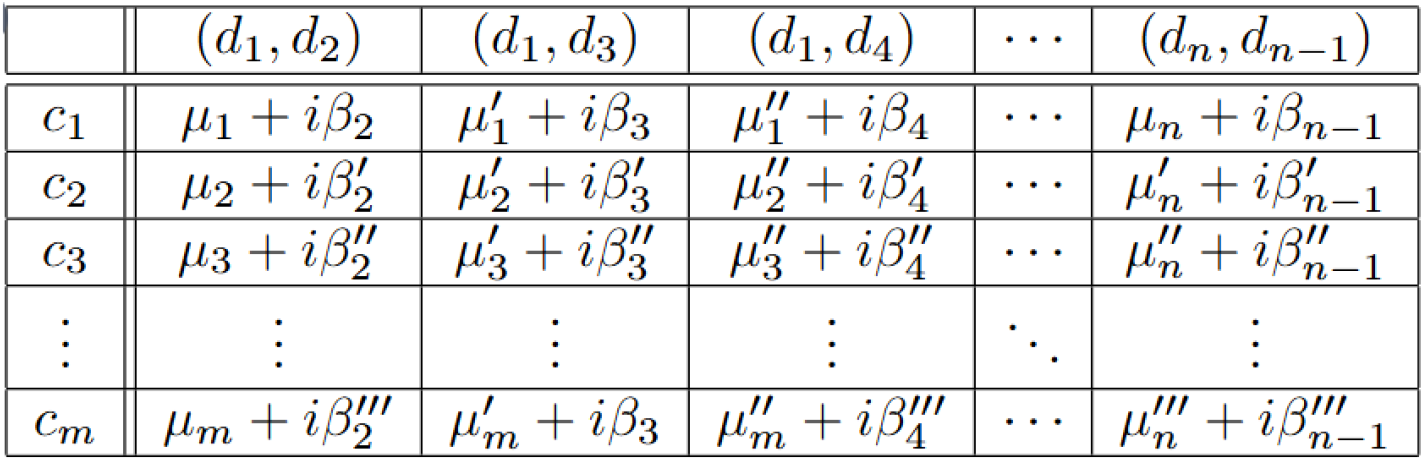
Schematic of the final processed matrix M, illustrating the drug-cell line relationships with complex entries.

#### 2.2.4 Step 4: Matrix Factorization Method

Let *CD* ∈ ℂ^*m×t*^ be the complex-valued matrix representing the interaction between *m* cell lines and *t* drug pairs. Our goal is to approximate *CD* using low-rank matrix factorization, i.e.,

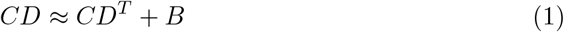

where:

- *C* ∈ ℂ^*m×h*^ is the complex-valued latent embedding matrix for cell lines,
- *D* ∈ ℂ^*t×h*^ is the complex-valued latent embedding matrix for drug pairs,
- *B* ∈ ℂ^*m×t*^ is a bias matrix,
- *h* ≪ min(*m, t*) is the dimensionality of the latent space.

The approximation is optimized by minimizing the following loss function based on the *L*_2_ norm:

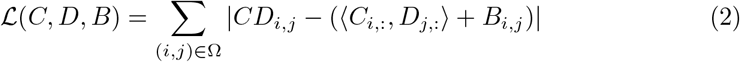

where Ω denotes the set of observed (non-zero) entries, and ⟨· ·⟩, denotes the complex inner product. Each matrix entry is approximated as:

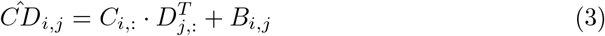

To optimize *C, D*, and *B*, we employ gradient descent using Wirtinger derivatives, which allow for differentiating real-valued loss functions with respect to complex variables. Specifically, for any complex variable *z* = *x* + *iy*, the gradients are computed as:

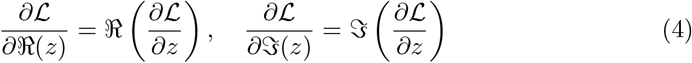

The parameters are updated iteratively as follows:

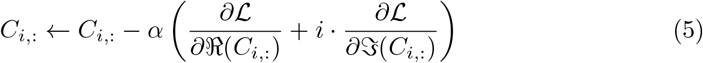

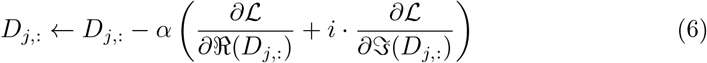

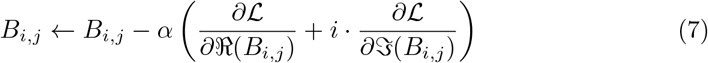

We also include an *L*_2_ regularization term on the latent matrices to avoid overfitting:

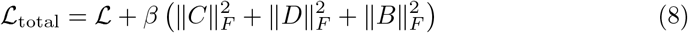

The algorithm is trained for a fixed number of epochs or until convergence.

This approach extends traditional matrix factorization methods [41, 42] into the complex domain. We adopt the Wirtinger calculus framework [43] to ensure valid and stable optimization for complex-valued parameters.

#### 2.2.5 Step5: Predict concentration values for *c, d*, and *d*^**′**^

After training the matrix factorization model, we obtain an embedding vector *C*_*c*_ for each cell line c and an embedding vector *D*_(*d,d*′_) for each drug pair (*d, d*^′^). Let the concentration values of the drug pair (*d, d*^′^) that inhibit the growth rate of cell line c by 50% are initially unknown. To estimate these concentration values, we calculate the dot product of *C*_*c*_ and *D*_(*d,d*′_). Following this calculation, we add the bias term *B*_(*c*,(*d,d*′_)), where *B* is the bias matrix obtained in Step 4 of the method. The resulting complex number represents the *log*_2_-transformed concentration value due to the normalization applied to the dataset. To retrieve the actual concentration values, we exponentiate the result using base 2. The resulting complex number is in the form *µ*_*d*_ + *i* · *µ*_*d*′_, where *µ*_*d*_ represents the concentration of the first drug d and *µ*_*d*′_ represents the concentration of the second drug *d*^′^.

#### 2.2.6 Implementation Details

The implementation settings used across all experiments are summarized in Table 1. These hyperparameters were selected based on empirical tuning to ensure a balance between convergence speed and predictive performance.

**Table 1.** Summary of Implementation Details.

| Hyperparameter | Value / Description |
| --- | --- |
| Hidden dimension size $h$ | 50 |
| Initialization | Complex Xavier initialization |
| Optimizer | Stochastic Gradient Descent (SGD) |
| Learning rate $\alpha$ | 0.01 |
| Regularization coefficient $\beta$ | 0.001 |
| Max number of epochs | 500 |
| Early stopping criteria | Relative loss change $< 10^{-5}$ |
| Random seed | Fixed across all runs for reproducibility |

## 3 Results

In this section, we evaluated our method on three publicly available datasets: O’Neil, NCI60-ALMANAC, and AZ-Dream. To ensure robust and reliable evaluation, we applied 5-fold cross-validation on all datasets. In 5-fold cross-validation, the dataset is divided into five equal-sized subsets (or folds). Each fold is used as a testing set exactly once, while the remaining four folds are used as the training set. The process is repeated five times, with each fold serving as the test set in turn. The final performance metrics are obtained by averaging the results across all five iterations, providing a comprehensive assessment of the model’s ability to generalize to unseen data.

This approach helps mitigate overfitting and reduces variability in the results by ensuring that all data points are used for both training and testing. Moreover, it provides insights into the stability and robustness of the method across different partitions of the data. Our evaluation consisted of two tasks regression and classification Tasks:

- **Regression Task**: This task aimed to predict the optimal dosages of drug pairs that inhibit 50% of cancer cell line growth. Predicted dosages were compared with the ground truth values provided in the datasets, and the accuracy was measured using the Mean Squared Error (MSE) and Mean Absolute Error (MAE). For complex-valued predictions, these metrics are defined as:

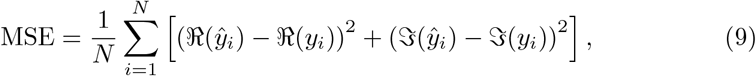

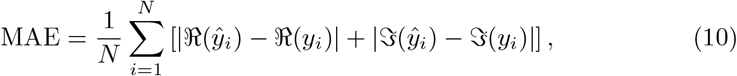

where ℜ (·) and ℑ (·) represent the real and imaginary parts of the predicted (*ŷ*_*i*_) and actual (*y*_*i*_) complex values, respectively.
- **Classification Task**: To further validate the performance of our model, we trans-

formed the regression task into a classification task. Each drug pair and cell line in our datasets is represented by constrained drug-response matrices, where known concentration values for each drug are defined. For example, in the O’Neil dataset, the drug-response matrix for a given drug pair and cell line is a 4 × 4 matrix, with four known concentrations for each drug. To evaluate classification performance we mapped the predicted continuous concentration values for each drug pair to their nearest neighbors within the set of known concentrations. After mapping, we assessed whether the predicted nearest concentration pair inhibited cancer cell line growth between 45% and 55%. To quantify the performance of our classification approach, we calculated the accuracy by measuring how often the predicted nearest concentration pair achieved an inhibition rate within the specified range of 45% to 55%. The accuracy metric is calculated as:

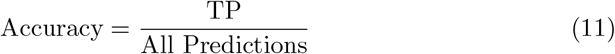

where TP (True Positives) denote the number of predicted concentration pairs that, after being mapped to their nearest neighbors, inhibit cancer cell line growth between 45% and 55%.

### 3.1 Results on the O’Neil Dataset

- **Regression Results**: We performed 5-fold cross-validation on 2,106 test data points per fold to predict the optimal concentrations of drug pairs for 50% growth inhibition. The regression results, summarized in Table 2, highlight the model’s predictive performance. After applying log normalization, concentration values were constrained to the range (-15, 9). The worst-case MSE occurs when the true concentration pair (9, 9) is predicted as (−15, −15), yielding an MSE of 1,250 and an MAE of 24 2. In contrast, across the 5 cross-validation folds, our model achieved significantly lower average MSE and MAE values of 6.27±0.13, and 2.21±0.02 respectively, where the values after the ± symbol represent the standard deviations. When converted back to the original scale, the average MSE and MAE were 35.99± 1.34 and 9.95± 0.58, both remaining far below the worst-case values. These results validate the model’s effectiveness in accurately predicting drug pair concentrations.
- **Classification Results**: We evaluated classification accuracy by measuring the proportion of correctly predicted drug-response pairs within the target inhibition range of [45%, 55%]. Our model achieved an accuracy of 30.46± 1.09, significantly surpassing the baseline random selection probability of at most 1/16 (Table 3). The baseline probability is derived from the 4×4 drug-response matrix structure used in the dataset, where each matrix element represents a unique combination of drug concentrations, resulting in 16 possible pairs in total.

**Table 2.** Performance Metrics For Regression Task Across Datasets.

| Dataset | MAE (Log) | MSE (Log) | MAE (Original) | MSE (Original) |
| --- | --- | --- | --- | --- |
| O’Neil | $2.21 \pm 0.02$ | $6.27 \pm 0.13$ | $9.95 \pm 0.58$ | $35.99 \pm 1.34$ |
| Almanac | $3.07 \pm 0.003$ | $12.26 \pm 0.20$ | $12.56 \pm 0.16$ | $23.25 \pm 0.37$ |
| AZ-Dream | $2.67 \pm 0.05$ | $9.51 \pm 0.32$ | $1.38 \pm 0.18$ | $6.12 \pm 1.024$ |

**Table 3.** Performance Metrics For Classification Task Across Datasets.

| Metric | O’Neil | ALMANAC | AZ-Dream |
| --- | --- | --- | --- |
| Accuracy (%) | $30.46 \pm 1.09$ | $38.6 \pm 1.12$ | $20.8 \pm 1.07$ |

### 3.2 Results on the ALMANAC Dataset

- **Regression Results**: We applied the same approaches used for the O’Neil dataset to the NCI-ALMANAC dataset. Specifically, we performed 5-fold cross-validation, with each fold containing 1,800 test data points. Unlike the O’Neil dataset, where the drug-response matrices were 4×4, the NCI-ALMANAC dataset uses 3×3 drug-response matrices. Each matrix represents 9 possible combinations of drug concentrations for a given drug pair and cell line. Log normalization constrained the concentration values to the range (-18, 7). In the worst-case scenario, where the true concentration pair (7, 7) is predicted as (−18, −18),the maximum MSE is 2,888, and the maximum MAE is 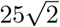. In contrast, across the 5 cross-validation folds, our model demonstrated significantly lower errors, achieving an average normalized MSE of 12.26±0.20 and MAE of 3.07±0.003. When converted back to the original scale, the average MSE and MAE were 23.25 ± 0.37 and 12.56 ± 0.16, respectively. (Table 2)
- **Classification Results**: We applied a similar approach as used for the O’Neil dataset to evaluate classification performance on the NCI-ALMANAC dataset. Specifically, we transformed the regression task into a classification task by mapping the predicted continuous drug concentrations to their nearest neighbors within the constrained 3×3 drug-response matrix of the ALMANAC dataset. As described for the O’Neil dataset this results in a random guess having a baseline probability of at most 1/9 for correctly identifying an effective concentration pair. Our model achieved an accuracy of 38.6± 1.12 (Table 3), which is 3.6 times higher than the baseline probability.

### 3.3 Results on the AZ-Dream Dataset

- **Regression Results**: We applied a similar approach as used for the O’Neil and NCI-ALMANAC datasets to the AZ-Dream dataset. Specifically, we performed 5fold cross-validation, with each fold containing 1,096 test data points, to predict the optimal concentrations of drug pairs for achieving 50% growth inhibition. The regression results are summarized in Table 2. After applying log normalization, concentration values were constrained to the range [− 16, 6]. In the worst-case scenario, where the true concentration pair (-16,-16) is predicted as (6,6), the maximum Mean Squared Error (MSE) and Mean Absolute Error (MAE) values were calculated as 968 and 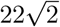, respectively. However, across the 5 cross-validation folds, our model demonstrated significantly lower average errors, achieving a normalized MSE of 9.51±0.19 and MAE of 2.67±0.04, where the values following the ± symbol represent standard deviations. When predictions were converted back to the original scale, the model exhibited similarly robust performance, achieving an average MSE of 6.12±0.27 and MAE of 1.38±0.08, both substantially below the worst-case scenario values.
- **Classification Results**: We also applied a similar classification approach as used for the O’Neil and NCI-ALMANAC datasets to evaluate the AZ-Dream dataset. The regression task was reformulated as a classification problem to identify drugresponse pairs within the target inhibition range of [45%, 55%]. Predicted continuous concentrations were mapped to the nearest neighbors within the 5×5 drug-response grid of the AZ-Dream dataset. The baseline probability for random selection was at most 1/25, derived from the 5×5 grid structure. In contrast, our model achieved an accuracy of 0.20±0.01 (Table 3), which is five times higher than the baseline probability.

These results highlight the robustness and reliability of our approach in identifying effective drug pair concentrations that inhibit cell growth within the desired range, further demonstrating the generalizability of our method across diverse datasets.

### 3.4 Comparing Complex Matrix Factorization with Alternative Machine Learning Methods

A key objective of this study is to assess the effectiveness of our matrix factorization-based approach compared to alternative machine learning methods. One possible alternative is to treat the problem as a multi-output regression task, where a model directly predicts the optimal drug concentrations, *µ*_*i*_ and *µ*_*j*_, based on a common set of input features. This approach differs from our complex matrix factorization method, which operates on a matrix with complex-valued entries. Instead of directly predicting drug concentrations, matrix factorization decomposes this complex-valued matrix to learn latent drug interaction patterns, enabling more accurate and generalizable predictions. To investigate this, we implemented a regression-based pipeline inspired by ComboFM. The data preprocessing steps remained consistent with ComboFM; however, instead of using factorization to infer drug interactions, we reformulated the problem as a supervised regression task. Here, the target outputs were the concentrations of *d*_1_ and *d*_2_, while the inhibition rate was included as an additional feature.

We also evaluated the following baseline models:

- ElasticNet, a linear regression method with L1 and L2 regularization.
- Random Forest, an ensemble learning approach that builds multiple decision trees.
- Multilayer Perceptron (MLP), a four-layer neural network trained to predict drug concentrations.

For ElasticNet, we used a regularization ratio of 0.5 (equal weight for L1 and L2 penalties) and performed grid search over alpha values in the range [10^−4^, 10^−1^]. For the Random Forest model, we used 100 trees with a maximum depth of 20 and employed mean squared error (MSE) as the splitting criterion. Hyperparameters were tuned using 5-fold cross-validation on the training data. The MLP consisted of four fully connected layers with 128, 64, 32, and 16 hidden units, respectively, each followed by a ReLU activation. The model was trained using the Adam optimizer with a learning rate of 0.001 and early stopping based on validation loss. Dropout with a rate of 0.2 was applied to mitigate overfitting.

All methods, including our complex matrix factorization model, were tested on three independent datasets—O’Neil, ALMANAC, and AZ-Dream—under identical experimental conditions. We used the same 5-fold cross-validation strategy described earlier, ensuring that all models were trained and evaluated using the same data splits. Table 4 presents MAE and MSE for each approach in the three datasets. The results clearly demonstrate that the complex matrix factorization-based method outperforms the alternative machine learning models, achieving consistently lower error rates. While the Random Forest model performed better than ElasticNet and MLP, it still lagged behind our method, particularly on the ALMANAC and AZ-Dream datasets. These findings suggest that performing matrix factorization on a complex-valued interaction matrix provides a more effective way to capture latent drug interactions, leading to more accurate dosage predictions than direct regression-based approaches.

**Table 4.**
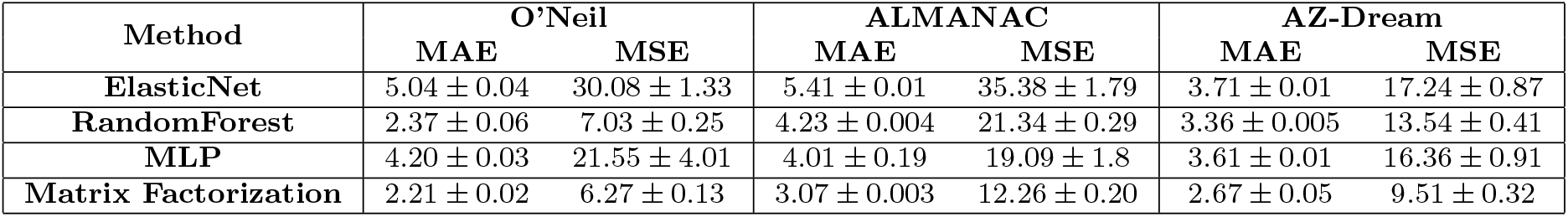
Performance comparison of our complex matrix factorization approach against alternative machine learning models (ElasticNet, Random Forest, and MLP) on three benchmark datasets (O’Neil, ALMANAC, and AZ-Dream). Metrics reported are Mean Absolute Error (MAE) and Mean Squared Error (MSE), averaged over five-fold cross-validation.

| Method | O’Neil |  | ALMANAC |  | AZ-Dream |  |
| --- | --- | --- | --- | --- | --- | --- |
|  | MAE | MSE | MAE | MSE | MAE | MSE |
| **ElasticNet** | $5.04 \pm 0.04$ | $30.08 \pm 1.33$ | $5.41 \pm 0.01$ | $35.38 \pm 1.79$ | $3.71 \pm 0.01$ | $17.24 \pm 0.87$ |
| **RandomForest** | $2.37 \pm 0.06$ | $7.03 \pm 0.25$ | $4.23 \pm 0.004$ | $21.34 \pm 0.29$ | $3.36 \pm 0.005$ | $13.54 \pm 0.41$ |
| **MLP** | $4.20 \pm 0.03$ | $21.55 \pm 4.01$ | $4.01 \pm 0.19$ | $19.09 \pm 1.8$ | $3.61 \pm 0.01$ | $16.36 \pm 0.91$ |
| **Matrix Factorization** | $2.21 \pm 0.02$ | $6.27 \pm 0.13$ | $3.07 \pm 0.003$ | $12.26 \pm 0.20$ | $2.67 \pm 0.05$ | $9.51 \pm 0.32$ |

As an alternative model architectures we also considered the possibility of replacing matrix factorization component with more complex deep learning architectures, such as Transformer-based models and other attention-based or graph-based neural networks. However, the available data was not sufficient to effectively train these high-capacity models. Despite experimenting with several configurations, their performance was significantly worse than our factorization-based method due to overfitting and lack of convergence. As a result, we do not include these models in the main comparison table, since their poor results were largely due to data limitations rather than methodological inferiority. Instead, we focused on more stable and reproducible baselines that perform reliably under low-data regimes.

### 3.5 Evaluating Model Predictions Using ComboFM and ComboLTR

While existing methods such as ComboFM and ComboLTR do not directly tackle the problem of predicting the exact concentrations of two drugs required to inhibit the growth rate of a given cell line by 50%, they address related challenges. Specifically, these methods take a different approach by predicting the inhibition rate of a given cell line based on a given drug pair and their concentrations. Since, to the best of our knowledge, no prior research has specifically focused on dosage prediction for a predefined inhibition rate, we leveraged these models as a validation tool rather than for direct comparison. To evaluate our results using these methods, we applied a validation strategy across the O’Neil, NCI-ALMANAC, and AZ-Dream datasets. We trained ComboFM and ComboLTR using the same train-test splits as our model to ensure consistency. Then, we tested how well our predicted drug concentrations integrate with these frameworks by evaluating two different scenarios:

1. Using Real Drug Concentration Values: The actual drug concentrations of the training sets were provided as input to ComboFM and ComboLTR.
2. Using Predicted Drug Concentrations: The drug concentrations predicted by our model were used as inputs to ComboFM and ComboLTR.

The accuracy results for both methods under these two scenarios are presented in Table 5. On the O’Neil dataset, ComboFM achieved an accuracy of 34.8% ± 0.01 when using real concentrations and 33.1% ± 0.01 when using our predicted concentrations, while ComboLTR achieved 31.7% ± 0.01 and 29.9% ± 0.01, respectively. On the ALMANAC dataset, ComboFM reached 25.8% ± 0.01 with real concentrations and 24.9% ± 0.01 with predicted concentrations, while ComboLTR attained 21.8% ± 0.01 and 20.6% ± 0.01, respectively. For the AZ-Dream dataset, ComboFM achieved an accuracy of 30.8% ± 0.0001 using real concentrations and 29.7% ± 0.05 with predicted concentrations, while ComboLTR achieved 26.4% ± 0.01 and 25.7% ± 0.04, respectively. These results indicate that, while the accuracy decreases when using predicted concentrations, the overall performance remains stable. Notably, ComboLTR maintained its ranking capability, suggesting that our model’s predictions align well with real data. This supports the use of our approach for downstream tasks, such as ranking drug combinations based on effectiveness. By leveraging ComboFM and ComboLTR as validation tools, we demonstrate that our predicted drug concentrations are sufficiently accurate to support drug combination analysis and precision oncology applications.

**Table 5.** Comparison of accuracy for ComboFM and ComboLTR using real vs. predicted drug concentrations on the O’Neil and ALMANAC datasets. In this table, RC and PC denote real concentrations and predicted concentrations, respectively.

| Method | O’Neil |  | ALMANAC |  | AZ-Dream |  |
| --- | --- | --- | --- | --- | --- | --- |
|  | RC | PC | RC | PC | RC | PC |
| **ComboFM** | $34.8 \pm 0.01$ | $33.1 \pm 0.01$ | $25.8 \pm 0.01$ | $24.9 \pm 0.01$ | $30.8 \pm 0.0001$ | $29.7 \pm 0.05$ |
| **ComboLTR** | $31.7 \pm 0.01$ | $29.9 \pm 0.01$ | $21.8 \pm 0.01$ | $20.6 \pm 0.01$ | $26.4 \pm 0.01$ | $25.7 \pm 0.04$ |

### 3.6 Robustness and Invariance to Drug Pair Ordering

In biological systems, the interaction between two drugs should not depend on their order in the dataset representation. To ensure that our model maintains this invariance, we assessed its robustness by evaluating the impact of drug pair reordering. This property is essential, as the designation of “first” and “second” drugs in a pair is arbitrary from a biological perspective. To test this, we randomly swapped the order of drugs in 50% of the dataset and trained our model using five-fold cross-validation on the O’Neil, ALMANAC, and AZ-Dream datasets. The same data splits were used for both the original and modified datasets to ensure a fair comparison. As summarized in Table 6, our model exhibited consistent performance across all key metrics, demonstrating its robustness to drug pair reordering. For example, on the O’Neil dataset, the MAE remained nearly identical between the original (2.21±0.02) and modified (2.20±0.02) datasets, with accuracy also showing minimal variation (30.42±1.14 vs. 30.46±1.09). Similar trends were observed for ALMANAC and AZ-Dream, where the differences in MAE, MSE, and accuracy between the original and reordered datasets were negligible. These results confirm that our model is invariant to drug pair ordering, ensuring its reliability in real-world applications where drug orders are not predefined. The observed stability across datasets further reinforces the robustness of our approach for modeling drug interactions in diverse settings.

**Table 6.** Performance metrics of the model on original and modified shuffled datasets for O’Neil, ALMANAC, and AZ-Dream.

| Dataset | Metric | Original Data | Modified Data |
| --- | --- | --- | --- |
| O’Neil | MAE | $2.21 \pm 0.02$ | $2.20 \pm 0.02$ |
| | MSE | $6.27 \pm 0.13$ | $6.27 \pm 0.12$ |
| | Accuracy | $30.42 \pm 1.14$ | $30.46 \pm 1.09$ |
| ALMANAC | MAE | $3.07 \pm 0.003$ | $3.06 \pm 0.003$ |
| | MSE | $12.26 \pm 0.20$ | $12.25 \pm 0.18$ |
| | Accuracy | $38.6 \pm 1.12$ | $38.7 \pm 0.008$ |
| AZ-Dream | MAE | $2.67 \pm 0.05$ | $2.67 \pm 0.06$ |
| | MSE | $9.51 \pm 0.32$ | $9.50 \pm 0.30$ |
| | Accuracy | $20.8 \pm 1.07$ | $20.9 \pm 1.05$ |

### 3.7 Experimental Prediction and Validation of Drug Pair Doses Using MTT Assay

In drug combination research, experimental datasets often suffer from incomplete coverage due to the high cost and time required for preclinical testing. As a result, many drug pair–cell line combinations lack experimental evidence for achieving specific inhibition thresholds, such as 50% growth inhibition. These gaps hinder the identification of effective drug combinations and limit progress in precision oncology. To overcome this challenge, we developed a two-phase strategy that integrates predictive modeling with experimental validation. Our model was trained on the complete available dataset to learn patterns in drug response and then used to impute missing values in drug–response matrices; specifically, combinations lacking IC_50_ data. To enhance reliability, we focused on drug pair–cell line combinations where the model demonstrated high prediction accuracy during 5-fold cross-validation. This targeted approach allowed us to prioritize high-confidence combinations for experimental validation, reducing the burden of exhaustive laboratory testing. Based on the model’s highconfidence predictions, the following drug pair–cell line combinations were selected from the ALMANAC and O’Neil datasets for experimental validation:

**From the ALMANAC dataset:**

– Ifosfamide and Gefitinib on OVCAR-3
– Ifosfamide and Gefitinib on HL-60
– Ifosfamide and Gefitinib on ACHN
– Ifosfamide and Gefitinib on HCT-116

**From the O’Neil dataset:**

– Gemcitabine and Sorafenib on HCT-116

#### 3.7.1 Experimental Validation Using MTT Assay

To test the validity of the predicted IC_50_ values, we conducted in vitro experiments using MTT assays on selected drug pair–cell line combinations. Four human cancer cell lines were selected for experimental validation:

- HCT-116 (colon cancer)
- OVCAR-3 (ovarian cancer)
- HL-60 (leukemia)
- ACHN (kidney cancer)

The details are demonstrated in Table 7.

**Table 7.** Predicted Drug Combinations and Applied Doses.

| Sample | Cell Line | Tissue Type | Drug 1 | Dose 1 (mg) | Drug 2 | Dose 2 (mg) |
| --- | --- | --- | --- | --- | --- | --- |
| 1 | OVCAR-3 | Ovary | Ifosfamide | 1.43 | Gefitinib | 2.337 |
| 2 | HL-60 | Hematopoietic | Ifosfamide | 1.214 | Gefitinib | 2.502 |
| 3 | ACHN | Kidney | Ifosfamide | 1.268 | Gefitinib | 5.270 |
| 4 | HCT-116 | Colon | Ifosfamide | 1.399 | Gefitinib | 2.864 |
| 5 | HCT-116 | Colon | Gemcitabine | $1.118 \times 10^{-3}$ | Sorafenib | 3.448 |

All cell lines were obtained from the National Cell Bank of Iran. Cells were cultured in RPMI-1640 medium (Gibco), supplemented with 10% fetal bovine serum (FBS) and penicillin-streptomycin (100 U/mL and 100 µg/mL, respectively). Cultures were maintained at 37 °C in a humidified incubator with 5% CO_2_ and passaged using trypsin-EDTA.

Cell viability was assessed using the MTT assay (Sigma-Aldrich). Cells were seeded in 96-well plates at a density of 1.4 × 10^4^ cells/well in 200 µL RPMI medium. After 24 hours, they were treated with predicted IC_50_ drug combinations and incubated for an additional 24 hours. Each condition was tested in technical triplicates and repeated across three independent biological replicates.

- Negative control (untreated)
- Vehicle control (DMSO-treated)
- Positive control (known cytotoxic drugs)

After treatment, the following *MTT* procedures apply:

1. Remove media and add 100 µL of MTT solution (0.5 mg/mL in PBS).
2. Incubate for 4 hours at 37 °C.
3. Remove MTT solution and add 100 µL of DMSO.
4. Measure absorbance at 570 nm using an ELISA reader.

The average absorbance in untreated control wells was 0.704, representing 100% viability. All drug-treated conditions resulted in a marked reduction in absorbance, confirming the cytotoxic effects of the predicted doses and aligning closely with IC_50_ expectations. The results were consistent across replicates. Slight variability observed in samples treated on OVCAR-3 and HL-60 may reflect cell line–specific sensitivities. No significant outliers were detected, indicating high reproducibility of the experimental protocol (Table 8).

**Table 8.** Observed Viability After Treatment.

| Sample | Cell Line | Drug Pair | Viability (%) | Standard Deviation (%) |
| --- | --- | --- | --- | --- |
| 1 | OVCAR-3 | Ifosfamide + Gefitinib | 46 | $\pm 10.05$ |
| 2 | HL-60 | Ifosfamide + Gefitinib | 51 | $\pm 8.23$ |
| 3 | ACHN | Ifosfamide + Gefitinib | 49 | $\pm 2.78$ |
| 4 | HCT-116 | Ifosfamide + Gefitinib | 48 | $\pm 1.56$ |
| 5 | HCT-116 | Gemcitabine + Sorafenib | 49 | $\pm 1.04$ |

These findings confirm that the predicted doses effectively induced approximately 50% reduction in cell viability, validating the model’s accuracy across multiple cancer cell lines and drug combinations. This outcome supports the broader utility of computational dose prediction in preclinical settings and underscores its potential to streamline experimental design in drug discovery and precision oncology.

## 4 Discussion

Combination therapy has emerged as a key strategy in the treatment of complex diseases such as cancer. By simultaneously targeting multiple pathways, drug combinations can improve therapeutic efficacy, reduce the likelihood of resistance, and minimize side effects. However, one of the central challenges in designing combination therapies lies in determining the precise concentrations of drugs needed to achieve specific biological outcomes, particularly the half-maximal inhibitory concentration (IC_50_), which reflects the dosage required to inhibit 50% cancer cell growth. While numerous computational models have been proposed to predict the inhibition rate or classify drug interactions as synergistic, antagonistic, or additive, there has been a notable absence of methods capable of predicting the exact drug concentrations required to reach a given inhibition threshold. Most existing approaches address the forward problem: predicting the effect given the drug and dose, whereas our work introduces a framework for the inverse problem: predicting the dose given a desired effect. To address this gap, we developed ComplexMatrixComb, a novel matrix factorization-based model that represents drug pair concentrations as complex numbers. This formulation captures the interdependent effects of two drugs using real- and imaginary-valued entries. By leveraging latent patterns in cell line–drug interactions, this model enables the prediction of optimal drug pair concentrations required to achieve IC_50_ in a specific cancer cell line. Our results demonstrate that this method significantly outperforms random selection and several established machine learning models across three benchmark datasets, e.i. O’Neil, ALMANAC, and AZ-Dream. In classification terms, while random chance offers an average accuracy of 1/9, 1/16 or 1/25 (depending on matrix size), our model consistently achieved much higher predictive accuracy, with results exceeding 20–38% across datasets. Furthermore, we validated our predictions by incorporating them into ComboFM and ComboLTR, two models designed to predict inhibition given a drug pair and concentration. When fed with concentrations predicted by ComplexMatrixComb, both models produced inhibition rates comparable to those using real concentrations, reinforcing the biological plausibility of our results. We also confirmed our predictions experimentally. Using MTT assays, we tested five drug pair–cell line combinations suggested by the model. All predicted concentration pairs resulted in viability near 50%, validating the accuracy and real-world applicability of our approach. The consistency of experimental outcomes across diverse cancer cell lines supports the robustness and translational potential of the method.

Importantly, ComplexMatrixComb is not limited to IC_50_ prediction. The framework can be extended to predict concentrations for any desired inhibition rate (e.g., IC70, IC90) by modifying the selection criteria during training. This flexibility broadens its utility across various stages of drug development and therapeutic planning. Another strength of our approach is its invariance to drug order, an important property, given that drug combinations are inherently symmetric. Our robustness analysis demonstrated stable performance when the order of drugs was randomized, further supporting the reliability of the model. Additionally, the model’s design allows it to generalize across datasets with different experimental resolutions and concentration matrices.

Despite its strengths, the model has some limitations. The use of complex matrix 20 factorization, while powerful, introduces interpretability challenges. Future work may incorporate biological priors or explainable AI techniques to enhance interpretability. Moreover, like most models, ComplexMatrixComb is based on data availability; a small pair of drug-cell line may limit its predictive capacity. Expanding the model to include other biological features, such as gene expression or pathway activity, could further enhance performance. In conclusion, this work presents a novel and practical solution to a previously unaddressed challenge in computational drug combination research. By accurately predicting the doses of drug pairs required to inhibit cancer cell growth, ComplexMatrixComb offers a transformative tool to accelerate drug discovery and optimize combination therapies. Its ability to reduce experimental workload, improve preclinical efficiency, and support clinical decision making makes it a valuable asset in the ongoing pursuit of personalized cancer treatment.

## 5 Conclusion

Identifying the precise concentration of a drug needed to inhibit the growth of a specific cancer cell line is a challenging and labor intensive task. For biologists, this often involves conducting extensive dose–response experiments across multiple concentrations, which becomes even more complex and resource-intensive when dealing with drug combinations. Testing all possible pairs at various dose levels quickly escalates into a combinatorial problem, requiring significant time, experimental effort, and financial cost. Traditional approaches rely almost exclusively on in vitro assays and iterative testing to determine the effective dose of a single drug or a drug pair. Although these methods provide accurate results, they are not scalable for large-scale drug screening or personalized therapy design. Computational tools have emerged to predict drug efficacy and synergy, but few, if any, offer a direct solution to the problem of estimating the exact doses required to reach a specific inhibition threshold such as IC_50_. In this context, our proposed method, ComplexMatrixComb, offers a practical and efficient alternative. Predicting the concentrations of drug pairs that induce 50% growth inhibition, it provides a data-driven solution to a problem that has traditionally relied on exhaustive laboratory tests. This capability has the potential to significantly reduce the experimental workload in preclinical drug development, streamline the design of combination therapies, and support more personalized and cost-effective treatment planning. As the need for accurate, scalable drug combination strategies continues to grow, tools such as ComplexMatrixComb can play a pivotal role in transforming how researchers and clinicians approach dose optimization in oncology. Despite its advantages, ComplexMatrixComb has some limitations. First, its performance depends on the availability of high-quality dose–response matrices, which remain limited for many drug pairs and cancer types. Second, while effective, the use of complex-valued matrix factorization can pose challenges to interpretability for biologists. Third, the current model was mainly validated in in vitro datasets, and additional tests in patient-derived or clinical data will be required to confirm its translational potential. Finally, due to the relatively small size of available datasets, we relied on a matrix factorization-based approach, which is more suitable for low-data regimes. With larger and more diverse datasets in the future, more expressive deep learning architectures, such as Transformer-based or graph-based models, could potentially be used to replace the factorization component, offering improved flexibility and performance.

## 6 Data & Code Availability

The source code and datasets are available at: https://github.com/mohammad−abdollahi/ComplexMatrixComb.

